# GDockScore: a graph-based protein-protein docking scoring function

**DOI:** 10.1101/2022.12.02.518908

**Authors:** Matthew McFee, Philip M. Kim

## Abstract

Protein complexes play vital roles in a variety of biological processes such as mediating biochemical reactions, the immune response, and cell signalling, with three-dimensional structure specifying function. Computational docking methods provide a means to determine the interface between two complexed polypeptide chains without using time-consuming experimental techniques. The docking process requires the optimal solution to be selected with a scoring function. Here we propose a novel graph-based deep learning model that utilizes mathematical graph representations of proteins to learn a scoring function (GDockScore). GDockScore was pre-trained on docking outputs generated with the Protein Data Bank (PDB) biounits and the RosettaDock protocol, and then fine-tuned on HADDOCK decoys generated on the ZDOCK Protein Docking Benchmark. GDockScore performs similarly to the Rosetta scoring function on docking decoys generated using the RosettaDock protocol. Furthermore, state-of-the-art is achieved on the CAPRI score set, a challenging dataset for developing docking scoring functions. The model implementation is available at https://gitlab.com/mcfeemat/gdockscore.

## 1 Introduction

Protein-protein complexes play a vital role in biological processes such as mediating biochemical reactions, signal transduction, and the immune response (Marsh & Teichmann 2015, Rebsamen et al. 2013). The structures of these complexes are key to their biochemical function (Sowmya et al. 2015) and structural biologists have been using techniques such as X-ray crystallography, nuclear magnetic resonance spectroscopy, and cryogenic electron microscopy (Curry 2015) to uncover them. However,these techniques are often difficult and time consuming (Nanev 2020) spurring the development of the in silico methods to accomplish a variety of tasks involving protein structure (Vakser 2014, Sesterhenn et al. 2020, Chevalier et al. 2017, Maia et al. 2020). Protein-protein docking is a two staged process where the initial stage involves sampling potential interfaces from the vast complex conformational space, and the second stage involves scoring of potential conformations (decoys) such that near-native interfaces are assigned favourable scores. This stage of the docking process is dependent on scoring functions which assess the energetic favourability of the proposed decoys (Kastritis & Bonvin 2011, Moal et al. 2013).

Protein-protein docking involves sampling the immense complex conformational space with the goal of determining the native interface between two polypeptide chains of interest. The conformational sampling process can be divided into three distinct search strategies including exhaustive global searching, local shape feature matching, and randomized searching (Huang 2014). In exhaustive global searching one protein is held fix while the rotational and translational degrees of freedom are explored by the other protein, this computationally intensive process can be expedited by applying Fast Fourier Transform techniques (Katchalski-Katzir et al. 1992). In local feature mapping techniques, the structures of the proteins can be represented as a Connolly surface and local complementarity of shapes is found between the two surfaces (Kuntz et al. 1982) or through geometric hashing where local matches in shape predictors are used to determine complementarity (Schneidman-Duhovny et al. 2005). Finally, in randomized searching one protein is randomly placed around the other (often with some constraint for the placements). The placement can then be refined using stochastic processes as Markov Monte Carlo Chains. After creating a set of candidate docking decoys, the second stage of the docking process begins, here decoys are scored based on their biophysical favourability using a scoring function.

Scoring functions for protein-protein docking can be placed into four distinct categories including force fields, empirical functions, knowledge-based functions, and machine learning based functions (Huang et al. 2010). Force fields involve calculations of physical atomic interactions such as van der Waals interactions and electrostatics (Harrison et al. 2018). In empirical scoring functions, energy terms such as van der Waals and hydrogen bonds are summed using learned weighting parameters (Pason & Sotriffer 2016). Knowledge based scoring functions employ statistical potentials to score interactions (Gohlke et al. 2000). Finally, machine learning based scoring functions leverage the increasing availability of protein complex structural data to learn scoring functions that can potentially have millions of parameters without the feature engineering involved in many traditional development techniques.

Previously, classical machine learning techniques such as support vector machines (Kinnings et al. 2011, Basu & Wallner 2016) and random forests (Li et al. 2015, Zilian & Sotriffer 2013) were used to develop scoring functions. However, these techniques still require feature engineering and are less performant than deep learning models. In deep learning, two types of network architectures have been used prominently to develop scoring functions. Convolutional neural networks have been used by discretizing the protein structure and generating 3D feature grids that are then passed through 3D convolutional layers (Wang et al. 2019, Schneider et al. 2021, McNutt et al. 2021, Renaud et al. 2021). Secondly, graph convolutional networks have been used where proteins are represented as mathematical graphs and node information is shared through message passing to learn a meaningful graph embedding (Cao & Shen 2020, Strokach et al. 2020, Geng et al. 2019, Wang et al. 2021). The use of convolutional layers is problematic in that they typically learn local information across the protein representation and cannot handle long distance relationships. Although there has been success in the use of dilated convolutional layers to learn relationships across greater distances in sequence data (Yu & Koltun n.d., Fudenberg et al. 2020). Finally, these networks require voxelization of the interface of interest which may result in loss of important information (Wang et al. 2021).

Our proposed scoring function utilizes a protein graph representation that has previously been very effective in learning embeddings for protein design and peptide binding prediction (Abdin et al. 2021, Ingraham et al. n.d.). This type of representation resolves the issues in standard convolutional networks. The protein graph embedding technique was further extended to learn graph embeddings of proteins in complex, which was done by developing a method to exchange interface information bi-directionally between protein graphs during the learning process. We show the effectiveness of our model architecture by learning a model pre-trained on RosettaDock (Roy Burman et al. 2018) generated on the PDB biounits, and fine-tuned on HADDOCK decoys (Dominguez et al. 2003, Renaud et al. 2021) using the ZDOCK Protein Benchmark dataset (Vreven et al. 2015). Our model performs well on RosettaDock decoys and on a gold-standard protein-protein docking scoring function development set known as the CAPRI score set (Lensink & Wodak 2014).

## 2 Results

### 2.1 GDockScore is a novel architecture utilizing protein graph attention and bi-directional interface information exchange

The model architecture utilized in GDockScore builds upon the parallel embedding architecture utilized by Abdin et al. (Abdin et al. 2021) and protein graph attention developed by Ingraham et al. (Ingraham et al. n.d.). Details of the model input representation are available in the Methods section. The model architecture is as follows: after an initial embedding of input node and edge matrices with linear layers, the basic building block of the architecture includes a protein graph attention block (Ingraham et al. n.d.) that updates the node embedding for each protein residue. In these blocks the edge matrices pass through linear layers to generate key and value pairs, and the node features are used to generate query vectors. The attention operation is then performed as seen in the original presentation of transformers (Vaswani et al. 2017) with the modification that only the 30 nearest neighbours to each node in the graph are used in the update to improve computational efficiency. As the Ingraham et al. protein graph attention is designed to update a protein graph embedding in isolation, we further developed a means of exchanging interfacial information between the two protein graph embeddings known as the bi-directional exchange layer.

In the bi-directional exchange layer the rows in the node embedding of the protein graph corresponding to interfacial residues are extracted into a separate matrix, hereby referred to as the interface embedding matrix. This matrix is used to generate a query matrix, while key and value pairs are generated from a matrix consisting of edge vectors from the interfacial residues of the current graph embedding to the 10 nearest neighbors on the other protein chain in the complex. As in previous models (Ingraham et al. n.d., Abdin et al. 2021), the corresponding node vectors on the adjacent proteins are concatenated to the appropriate edge vectors. This concatenation allows each protein embedding to directly “see” the residues of the other protein via node features. The standard attention mechanism is then performed to generate an update to each of the interface residue embeddings. The interface residue embeddings in the node matrix for the entire protein are then replaced with these updated interfacial residue node embeddings. This process is then repeated in the other direction to update the other protein chain in the complex. This basic module is stacked multiple times to generate final protein embeddings for each protein which are then concatenated and pooled down to produce a final output which passes through the sigmoid function to generate a probability of being a near-native decoy. The overall model architecture is illustrated in Figure 1 and further details of model hyperparameter tuning are available in the Methods section.

**Figure 1:**
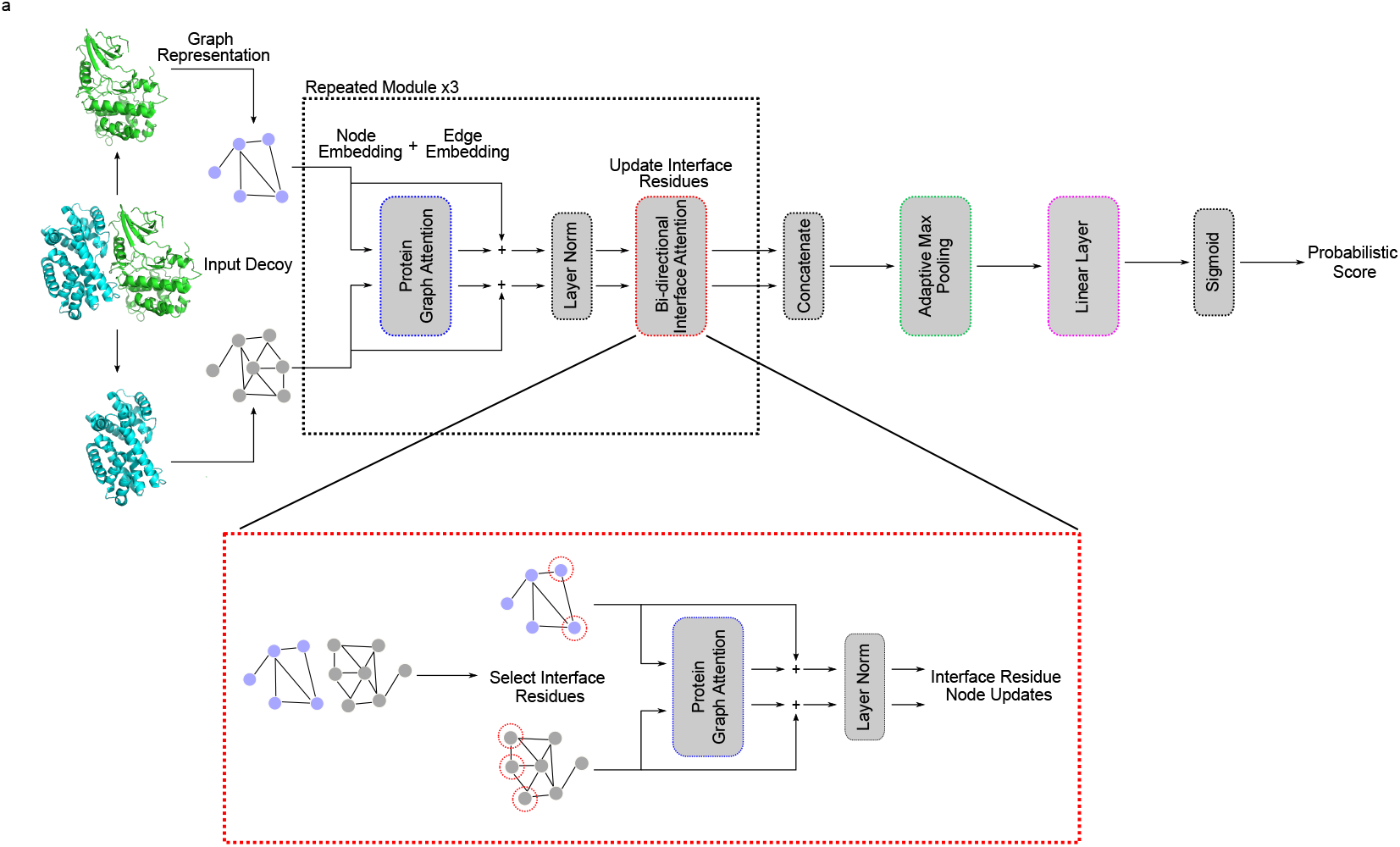
(A) The model architecture described in this paper. The inlaid image (dotted red line) illustrates the bi-directional exchange of information between protein graph embeddings. The protein depicted is the AB interface from PDB code 1f5q’s biological assembly 1.

### 2.2 Generation of a novel protein-protein docking pre-training dataset leveraging the PDB biounit files for model pre-training

Previously, protein-protein docking deep learning models were trained on gold-standard datasets such as the ZDOCK protein-protein docking benchmark (Hwang et al. 2010), and Dockground (Kundrotas et al. 2017). These datasets rely on decoys generated for a small number of targets and thus do not leverage the full complexity of structures and interfaces exhibited in the PDB. We generated a significantly more varied dataset for model pre-training by scanning all PDB biounit files for interacting chain pairs (snapshot March 2021).

To generate a positive dataset we extracted interacting chain pairs from the PDB biounits which are the proposed functional forms of protein complexes. This was done by searching for complexes that had at least 3 pairs of amino acids with an *α*-carbon to *α*-carbon distance of 6 Å or less. Not all of these were used as class balancing was conducted, and some structures failed to regenerate near-native decoys after docking with RosettaDock. The standard metric of CAPRI rank was used to indicate the quality of the decoys where 0 is incorrect, 1 is acceptable, 2 is medium quality, and 3 is high quality (Méndez et al. 2003). With these classifications depending on standard metrics such as interface root-mean-square deviation, and the fraction of contacts in the true interface recapitulated in the proposed decoy. The decoys used to train our model were generated by using the RosettaDock protocol (Roy Burman et al. 2018). For positive examples, the ligand protein was initialized to a position near the true interface with a small perturbation allowing for the ligand to re-dock to the receptor protein in a near-native configuration. To generate the negative dataset, the ligand protein initial position was completely randomized around the receptor protein and the receptor was also rotated randomly. The docking protocol was then carried out generating completely new interfaces with CAPRI ranks of 0 indicating significant difference from the native interface. This data generation process is depicted in Figure 2a.

**Figure 2:**
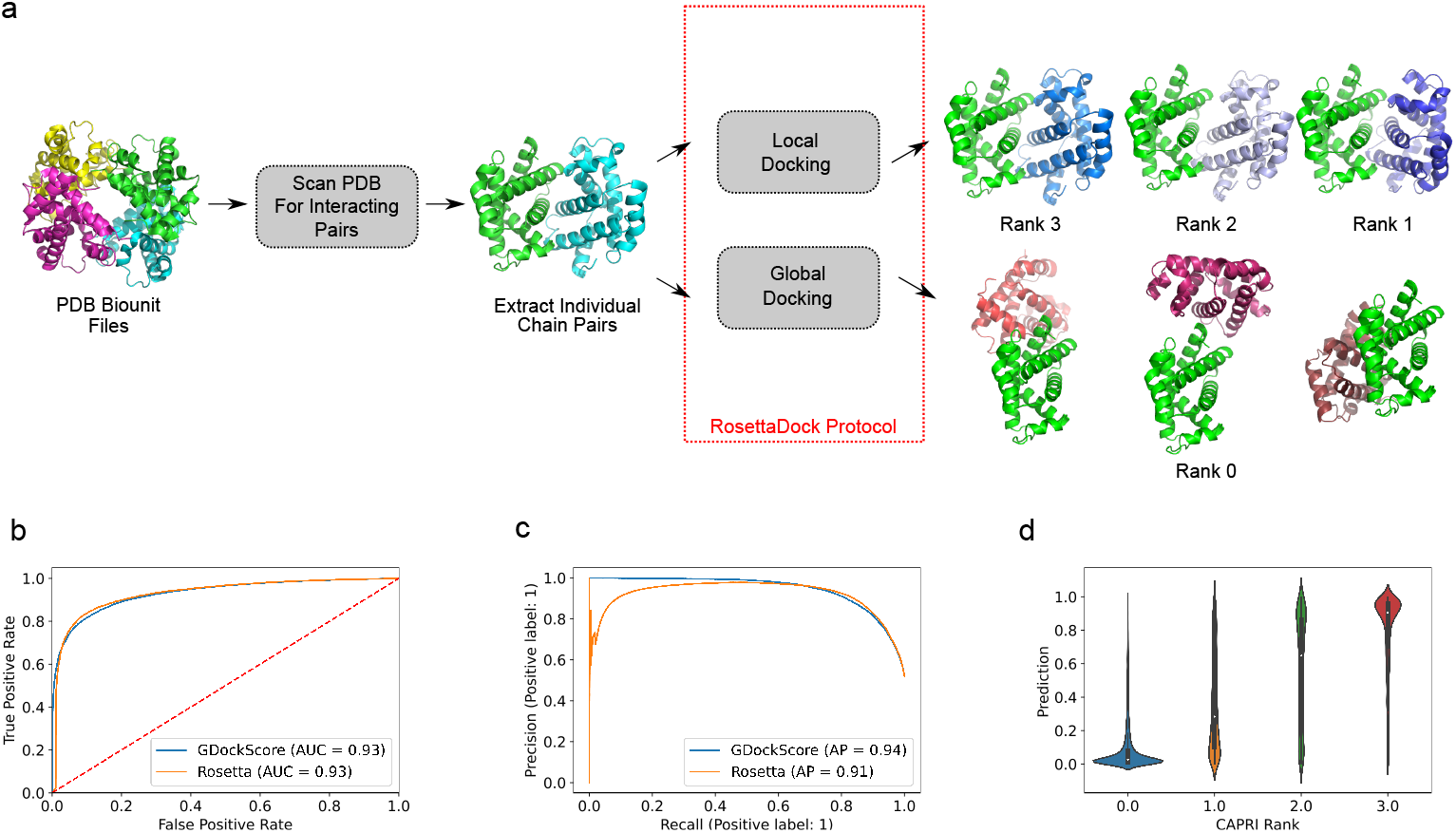
(a) Schematic illustration of the data generation process using the PDB biounit files used to create the model pre-training set. The biological assembly 1 for PDB code 1bab is illustrated.(b) The ROC with indicated AUC for our model in comparison to Rosetta’s interface score (I_sc). (c) The precision-recall curves for our model vs. I_sc with AP indicated. (d) Violin plot showing the distribution of predictions for each CAPRI category using our models output.

The above data generation protocol resulted in significantly more negative docks being generated in comparison to CAPRI 1, 2, and 3 positive examples. This is due to many chain pairs failing to generate CAPRI acceptable decoys even when initialized very close to the true interface and docked with RosettaDock as well as the algorithm generally being able to recover a solution very close to the true interface in the local docking protocol. To balance the data a subset of CAPRI 0, 1, 2, and 3 decoys were sampled such that the total number of positive examples equaled the negative examples. Due to the generation process, there were roughly an equal number of CAPRI rank 3 decoys to total number of rank 1 and 2. We kept this imbalance as we want to favour protein interfaces that are as close to the optimal interface as possible. To ensure no significant similarity between the training and validation sets, both sequence and structural analysis were performed. The MMseqs2 (Steinegger & Söding 2017) tool was used to cluster all unique protein sequences in the pre-training set based on similarity with a 30% cut-off point. Furthermore, to ensure that no remaining examples in the training, validation, and test sets had any similarity to our intended external test set (Lensink & Wodak 2014), any structure containing a chain clustered with the CAPRI score set members was removed. As an additional check, the US-align (Zhang et al. 2022) tool was used to calculate TM-scores (Zhang & Skolnick 2007) to find the structural similarities between the elements of the training, validation, and test sets and the targets of the CAPRI score set, with any example with a score >= 0.4 being removed as well, which should eliminate any remotely similar structures (Xu & Zhang 2010).

The standard RosettaDock protocol only adjusts side chain conformations at the interface and does not predict backbone conformational changes. Based on initial model analysis, we hypothesized could be supplemented with additional data. The DeepRank (Renaud et al. 2021) dataset includes HADDOCK (Dominguez et al. 2003) docking decoys generated using the ZDOCK Protein Docking Benchmark (BM5.5) (Vreven et al. 2015) data set. We exclusively selected the decoys that had backbone refinement performed at the interface (“it1” decoys) and those further refined with a short molecular dynamics simulation in water (“itw” decoys). In this set all decoys with an iRMSD ≤ 4 were considered positive examples. The pre-trained model parameters were then fine-tuned with this dataset.

### 2.3 GDockScore learns to perform well on predicting near-native from incorrect decoys generated by RosettaDock

First, we sought to assess the models performance on a test set consisting of decoys generated with our pre-training dataset generation scheme. The final AUC achieved by the trained model on the test set was 0.93 indicating that the model effectively learned to classify unseen protein-protein docking decoys with structures and sequences differing from those seen in the training set (Figure 2b). The fine-tuning on the HADDOCK decoys does not appear to significantly impact GDockScore’s ability to classify RosettaDock outputs. Furthermore, the average precision as determined by the sklearn package (Pedregosa et al. 2011) for the model was 0.94 (Figure 2c). We sought to directly compare the performance of the learned model to the classical Rosetta scoring function (Alford et al. 2017). For each decoy, the Rosetta interface score (I_sc) was used to classify the decoys as being near-native or incorrect. In comparison to our learned model, Rosetta achieved 0.93 AUC on the test set (Figure 2b). Thus, it appears that our model at least learns to recapitulate the Rosetta scoring function t. This suggests that fine-tuning on the HADDOCK decoys does not significantly reduce the models ability to score RosettaDock decoys. More interestingly, when comparing the average precision (AP), see equation 3 in Methods for details, our model outperforms Rosetta. Rosetta achieves an AP of 0.91 compared to our AP of 0.94. This indicates that our model typically achieves a greater precision in comparison to the standard Rosetta scoring function.

To better understand how the model was performing on individual CAPRI ranks, the model predictions for each rank were plotted as a violin plot (Figure 2d). The model appears to confidently predict CAPRI rank 3 and CAPRI rank 0 models as indicated by the majority of the violin plot’s density being focused at very high and very low scores. The CAPRI rank 3 models are very close to the native interface with interface root-mean-square deviation (iRMSD) values ≤ 1 and the CAPRI rank 0 decoys will have very large iRMSD values > 4. Intuitively, the model appears to struggle more on CAPRI rank 1 and 2 models which are deemed to be acceptable and medium quality. Therefore, it can be concluded that our model can confidently determine an almost exact interface from a completely incorrect one but it does struggle with lower quality models that are still somewhat near the true interface ie. CAPRI rank 1 decoys. This is logical, as the CAPRI rank 1 models can have iRMSDs as high as 4 which indicates a substantial deviation from the true interface and even small rotations and translations may result in unfavourable amino acid contacts being made at the decoy interface.

As a final check of the model in predicting high quality interfaces, we applied our model to the new, but related task of crystal and biological interface detection. Crystal interfaces are protein-protein interfaces that appear during the generation of protein crystals in X-ray crystallography experiments that are not biologically relevant. The MANY (Baskaran et al. 2014) and DC (Duarte et al. 2012) datasets of real biological and crystal interfaces were combined and scored by the model. The model achieved an AUC ROC of 0.93 and AP 0.92 on the precision-recall curve, indicating strong performance on differentiating crystal interfaces from biological interfaces even though the model was trained on protein-protein docking decoys (Supplementary Figure 1a,b). The model is more confident at classifying the biological interfaces than the crystal (Supplementary Figure 1c). These performance metrics are likely biased as there is sequence redundancy in the training set. However, this result is still promising in showing that the model architecture can easily be extended to similar tasks to protein-protein docking.

### 2.4 GDockScore outperforms state-of-the-art models on the challenging CAPRI score set

We sought to test our model on an externally generated gold-standard test set to verify its performance. We used the CAPRI score set (Lensink & Wodak 2014) a dataset for developing protein-protein docking scoring functions. This dataset consists of decoys generated for 13 unique chain pairs generated by 47 unique docking methods and is noted for being challenging. The version of the CAPRI score set used in this paper was the cleaned, and processed copy used in the development of iScore and DeepRank (Geng et al. 2019, Renaud et al. 2021). We benchmarked our model against DeepRank (Renaud et al. 2021) and GNN DOVE (Wang et al. 2021), two recent highly performant deep learning models.

We sought to compare our model against the others using the benchmark of AUC of the ROC, AP of the precision-recall curve, as well as comparing violin plots of score distributions for near-native vs. incorrect decoys. In terms the AUC of the ROC our model performs better than both DeepRank and GNN DOVE with an AUC of 0.86 compared to 0.72, and 0.54 respectively (Figure 3a). It is important to note that the initial model without any fine-tuning on the DeepRank dataset only achieves an AUC of 0.72, indicating that our fine-tuning set is providing significant useful information to GDockScore. Our model also achieves an improved AP in comparison to these two models, achieving 0.40 in comparison to 0.23 for DeepRank, and 0.23 for GNN DOVE (Figure 3b). The model with no fine-tuning achieves an AP of 0.19. To determine how well the models were differentiating the two classes of near-native and incorrect interfaces, a violin plot was generated (Figure 3c). GDockScore seems to do a slightly better job at applying lower scores to the incorrect decoys, where as DeepRank likes to score most decoys in the CAPRI score set highly. Our model also seems to achieve more density at high scores for the near-native class in comparison to DeepRank. In stark contrast, GNN DOVE appears to score all models with very low scores for both the near-native and incorrect classes. It is clear that for all state-of-the-art models, the CAPRI score set is still quite challenging, but GDockScore has achieved state-of-the-art performance.

**Figure 3:**
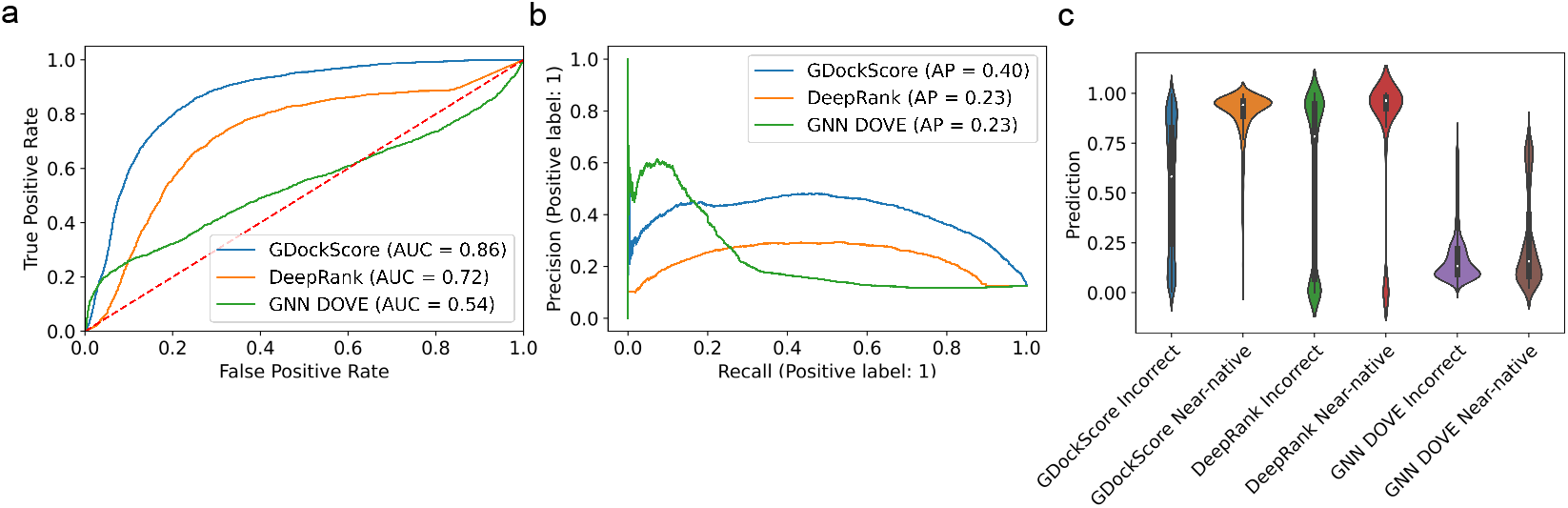
(a) The ROC for GDockScore, DeepRank, and GNN DOVE where AUCs are indicated in the inlayed figure legend and the red dotted line indicates luck. (b) The precision recall curves of our model, DeepRank, and GNN DOVE with AP indicated in the inlayed legend (c) Violin plot of each models predictions for near-native vs. incorrect models. Note: DeepRank predicts probability of the model being incorrect so we have taken 1 minus this probability so that each model output is in agreement.

Finally, we assessed the models performances on each individual CAPRI target by comparing the number of near-native structures in the top *N* ranked structures across a wide range of *N* (Figure 4). Our model performs comparatively to DeepRank and GNN DOVE across most targets. However, there are many cases where our model appears to outperform DeepRank and GNN DOVE. For example, T29, T32, T37, T40, T53, and T54. This is because our model is putting a larger number of near-native hits in the top N decoys at earlier values of N. This suggests that our model is, in general, assigning higher ranks to near-native decoys in the CAPRI score set which is the primary goal of a scoring model for tasks such as ranking docking decoys.

**Figure 4:**
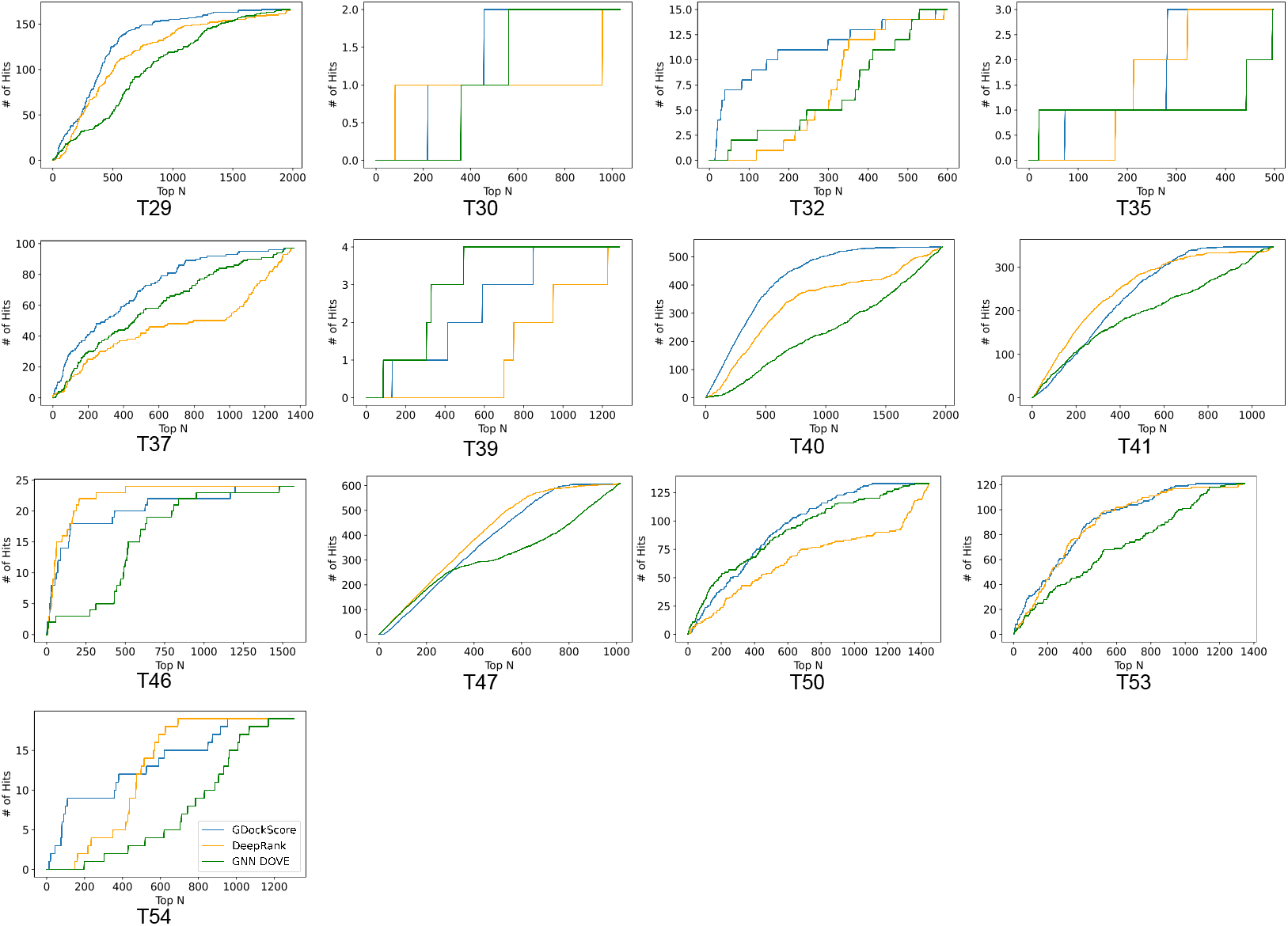
The number of near-native structures (# of Hits) in the Top *N* ranked structures for each of the CAPRI score set targets. Here we display the results of our model in blue, DeepRank in orange, and GNN DOVE in green.

## 3 Discussion

In this work we presented GDockScore, a deep learning based model that uses protein graph attention to learn a scoring model for protein-protein docking. We generated a new data set that leverages a substantial portion of the PDB biounits to pre-train GDockScore. We then fine-tuned on a small subset chains that underwent backbone refinement or a short molecular dynamics simulation. Our model effectively learned to classify decoys generated by the RosettaDock protocol as acceptable or better according to CAPRI standards, or incorrect. The model also shows promise in the task of distinguishing biological interfaces from crystal interfaces showing the potential applicability of GDockScore to new protein structure classification tasks.

After this, we sought to challenge our model on the CAPRI score set a difficult data set consisting of decoys from many different docking protocols. GDockScore shows better performance compared to other state-of-the-art deep learning models on the CAPRI score set. This includes better AUC ROC and AP of the precision-recall curve in comparison to two highly performant models, DeepRank and GNN DOVE. Upon inspection of number of hits (near-native decoys) in the top N structures, over a complete range of N, GDockScore out performed DeepRank and GNN DOVE in several instances as indicated by having more hits in smaller values of N. This is crucial as these scoring models are used to rank decoys and it is optimal to maximize the number of hits in the N selected best scoring structures.

There are a variety of ways in which this work can be extended. It is clear that all recent models find the CAPRI score set to be particularly challenging. Similarly to our model having a performance drop from Rosetta decoys to the CAPRI score set when only RosettaDock decoys were used, DeepRank performs much better on classifying the HADDOCK decoys. This drop in performance of models on the CAPRI score set when only one docking protocol is used to generate input data is likely due to the challenging nature of the chain pairs in the set, as well as the variety of algorithms used to generate the decoys. It may be possible to improve the current deep learning models by training on a data set generated with many different docking algorithms to reduce the likelihood that a model trained on decoys generated by a single docking algorithm is learning the individual idiosyncrasies of said algorithm.

Another limitation of our model is the fact that the data set we introduced extracted individual chain pairs from each PDB biounit. Therefore, the context of the chains interacting in a complex of greater than two chain pairs are lost. Model performance may be improved by training the model to embed complexes of more than two chains at the same time.

In summary, we have developed a new deep learning scoring function for classifying protein-protein docking decoys that performs strongly on decoys generated using the RosettaDock protocol, and the CAPRI score set, a challenging test set in protein-protein docking scoring.

## 4 Methods

### 4.1 Details of the Input Representation

The input features of the model graphs are based on work developed by Ingraham et al. (Ingraham et al. n.d.) and Abdin et al. (Abdin et al. 2021). To generate rotationally, and translationally invariant structural features, local coordinate systems are established at each *α*-carbon along the protein backbone allowing for vectors in the global reference frame to be transformed to each protein residues local coordinate system, providing rotational and translational invariance.

The node features of the graph include the one-hot-encoded amino acid identity, distances from the *α*-carbon to each heavy side chain atom raised to a radial basis, directional vectors from each *α*-carbon to each heavy side chain atom transformed to the local coordinate system, as well as backbone torsional angles embedded in the 3-torus. Our work builds upon Abdin et al. by extending the side chain representation information from a course centroid representation to a much more granular all heavy atom representation. Edge features are represented via *α*-carbon to *α*-carbon distances raised to a radial basis, directional vectors transformed to the local coordinate system, as well as quaternion representation of the dot product of the two coordinate systems at each *α*-carbon which captures information about relative coordinate system orientations. Information about relative distance in sequence was encoded in the way described in the original implementation (Ingraham et al. n.d.). Inter-chain edges contain the same information as edges within a protein graph excluding the encoding of relative distance in protein sequence.

### 4.2 Details of the Bi-directional Attention Block

In the bi-directional attention block there are two directions for the update. Suppose there are two proteins which will be referred to as A and B. The interface residue nodes of the matrix are selected using indexing. The interface residue update for protein A is as follows:

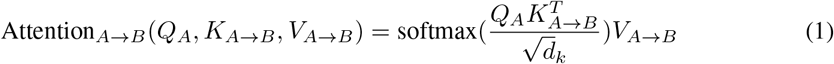

Where the query matrix *Q_A_* is a ℝ*^n×d_κ_^* matrix generated using a linear layer and the node embeddings of A’s interface residues. The key matrix *K*_*A*→*B*_ is ℝ*^n×d_κ_^* and the value matrix *V*_*A*→*B*_ is ℝ*^n×d_v_^*, with both being generated using the edge matrix which consists of edges from the interface residues of A to the 10 nearest neighbours on protein B. Here, the symbol → indicates that we are updating the protein on the left of the arrow with information from the protein on the right of the arrow. The updates for B are generated in a similar manner but instead extracting the interface residues of the node matrix of B and using an edge matrix consisting of edges from B to A. It is also crucial to note that the corresponding node features are concatenated onto the edge features allowing direct exchange of node information from one protein to the other. Thus, if there is an edge from node *A_i_* to node *B_j_*, where *i* and *j* are residue numbers in each protein, then the edge *e_ij_* will be concatenated to node embedding Bj before the key value pair generation. For more details of the generation of the query, key, and value matrices please consult Ingraham et al. (Ingraham et al. n.d.).

### 4.3 Model Hyperparameter Tuning and Training

Model hyperparameter tuning was performed by varying 8 hyperparameters including *d_model_* (the model embedding dimension), *d_inner_* (the dimension of the hidden layer in the feed forward layers), *d_k_*, *d_v_*, the dropout percentage, the number of repetitions of the exchange layer, the number of heads in each attention layer, and the learning rate. The final tuned parameters were *d_model_* = 16, number of heads = 3, number of layers = 3, *d_k_* = 16, *d_v_* = 32, and *d_inner_* = 16.

Training was performed using the Adam optimizer (Kingma & Ba 2014) with a learning rate of 1 * *e*^-4^ which was determined to be optimal via tuning. The loss function was binary cross-entropy as implemented by PyTorch (Paszke et al. 2019). Early stopping was implemented by monitoring the validation loss during training. The model typically converged at approximately 1 epoch of training. Fine-tuning was performed with a learning rate of 1 * e^-5^ and the model converged after a little over an epoch of training.

### 4.4 Definition of Average Precision

The average precision reported is from the implementation provided by the scikit-learn machine learning package (Pedregosa et al. 2011). The mathematical definition is

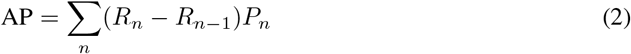

Here AP represents average precision, the summation is over the *n* thresholds, *R_n_* is the recall at the given threshold, *R*_*n*-1_ at the previous threshold, and the weighting term *P_n_* is the precision at the current threshold.

## 5 Acknowledgements

We would like to acknowledge Osama Abdin for his general feedback on the project, and the feature generation code that served as the basis for the code included in this paper. We’d also like to thank Sunyun Lee for reviewing some of our code.

## 6 Contributions

The model architecture was conceptualized by MM and PMK. Implementation, data generation/curation, and data analysis were performed by MM. Writing was performed by MM and PMK. Supervision and funding acquisition is attributed to PMK.

## 8 Data and Code Availability

The data set introduced in this paper is available at http://gdockscore.ccbr.proteinsolver.org. The CAPRI score set files used in this paper are available at https://data.sbgrid.org/dataset/684/. The DeepRank data set is available at https://data.sbgrid.org/dataset/843/. The DC and MANY data sets are available at https://www.eppic-web.org/ewui/#downloads. The code is available at https://gitlab.com/mcfeemat/gdockscore.

## A Supplementary Information

**Figure 1:**
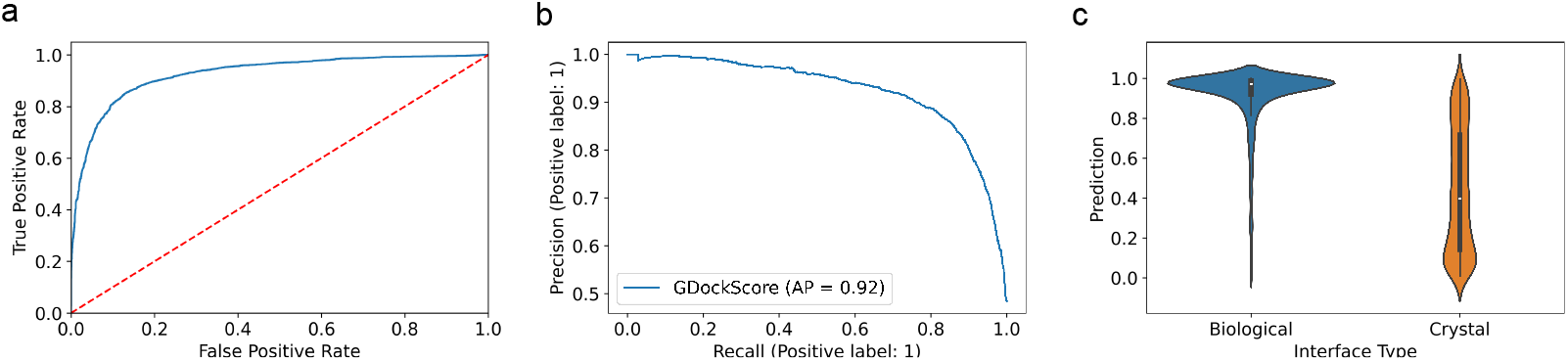
(a) The ROC of the model on the MANY + DC combined data set (AUC = 0.93). The dotted red line indicates pure luck. (b) The precision-recall curve of the model on the MANY + DC combined data set. The inlayed legend contains the AP. (c) Violin plot depicting the distribution of scores for each class of interface (biological vs. crystal).

